# Application of dielectric spectroscopy in the dynamically changing biological system – case of *Neocaridina davidi* shrimp starvation

**DOI:** 10.1101/2020.06.22.164517

**Authors:** Agnieszka Wlodarczyk, Patryk Wlodarczyk

**Affiliations:** Department of Molecular Biology and Genetics, Medical University of Silesia, in Katowice, ul. Medyków 18, 40-752 Katowice, Poland; Lukasiewicz Research Network – Institute of Non-Ferrous Metals, ul. Sowinskiego 5, 44-100 Gliwice, Poland

**Keywords:** Neocaridina davidi, dielectric spectroscopy, starvation

## Abstract

In this work, dielectric studies on *Neocaridina davidi* shrimps have been presented. The effect of starvation on dielectric properties such as conductivity or permittivity have been shown. It was found that the onset frequency of electrode polarization depends on starvation period, which is probably related to the cytoplasm viscosity. In the dielectric spectra of shrimps two relaxation processes have been identified i.e. α and β process. The α process is probably related to the counter ion polarization while β process to the mobility of macromolecules present in the body of a shrimp, mainly to the amount of lipids. It was also found that there is a difference in dielectric response between control group and the group regenerated after 14 day starvation period. Basing on dielectric response, one can conclude that the viscosity of cytoplasm of regenerated shrimps is higher and the cells are rich in the lipid droplets when compared to control group.

## 1. Introduction

The *Neocaridina davidi* shrimp (formerly *Neocaridina heteropoda)* is a freshwater crustacean originating in Taiwan. It is widespread all over the world, due to the fact that it is popular breeding species. This work concentrates on the changes during the starvation of shrimp that can be monitored by dielectric spectroscopy. The changes are observed globally on the entire body of shrimp. Starvation is a natural process that can occur in environment. Many invertebrates hibernate during overwintering, while they are active during spring, summer and autumn. Therefore, the overwintering is the natural period of starvation [1,2]. Until now, many works related to the physiological, biochemical and histological changes during starvation of crustaceans have been published [3–7]. The results presented in this works are directly related to the earlier works describing ultrastructural changes in the midgut of *N. davidi* during starvation [8–10]. In the cited works, the average time of starvation after which regeneration of shrimps is impossible has been obtained. This parameter i.e. point of no-return PNR_50_ has been estimated to 24.7 days. The shrimps can survive extremely long period of time without food due to the fact that adults have accumulated storage material in the form of lipids in the cytoplasm of the cells in hepatopancreas. During the starvation period, increasing amount of inactive mitochondria i.e. mitochondria with lower membrane potential was observed. Refeeding shrimps for 14 days after 14 days of starvation was not enough to restore mitochondria activity to the level of control group [8].

Other studies indicated that the intensity of autophagy and apoptosis has also increased, which may indicate that it is a part of survival strategy. Moreover, after 14 days of refeeding, the apoptosis level was found to be lower than in the control group [9]. Starvation is causing oxidative stress in living organisms. The significantly increased levels of MnSOD radicals and antioxidants in the midgut were observed during the experiment. The effect is strongest in the initial fasting period. Therefore, this stressor may affect the impairment of the protection system against free radicals [10].

During the stress induced by starvation, the composition of shrimps as well as the alteration of their physical parameters occur. While the described earlier methods allowed for monitoring the processes occurring on the ultrastructural level in certain organs, dielectric spectroscopy will measure whole body response and the macroscopic, physical properties. Dielectric spectroscopy is a method that can be used for measurements of biological systems, in order to obtain electric permittivity and conductivity in the broad range of frequencies [11]. In the dielectric spectrum of tissues or cell suspensions there are usually three dispersions. The first process in the lowest frequencies is α-relaxation, which nature is not fully understand but it is probably governed by so called counter-ion polarization of cell membranes. It is very hard to monitor α-relaxation as it is usually covered by the interfacial polarization of electrodes occurring at low frequencies. The second dispersion, named β can be caused by the relaxations of macromolecules present in the tissues. The fastest process that can be observed in the GHz region is a water relaxation within the cells.

The starvation is a stress factor which affects physiological processes in the living organisms. On the macroscale it should alter functioning ion pumps in the cells, which should be visible in the low frequencies of dielectric spectrum (10 mHz – 100 Hz). There is also possibility that in the higher frequencies, beta relaxation peak will be affected due to the changes in the composition of the body, especially decreased amount of lipids.

The main aim of this work is to study changes of dielectric signal during starvation of shrimps and to compare these results to the changes in the ultrastructure of studied earlier organs.

## 2. Materials and Methods

The shrimps were held in the separate 250 mL containers during starvation. The containers were filled with the water from the main tank. The breed water parameters were as follows:

- water overall hardness 7-10 °d,
- pH in the range 6.8-7.0
- temperature 21-24 °C

The 10% of water in containers was exchanged daily. Moreover containers were cleaned from excrements on the daily basis. Containers were held in shady place in order to avoid algae development. The overall amount of shrimps for the experiment was set to 25. Five shrimps for every stage were used for studies. In the experiment five stages were chosen i.e. control group, shrimps starved for 3 days, 7 days, 14 days and shrimps fed 14 days after 14 days of fasting.

Shrimps from all stages were measured by means of broadband dielectric spectroscopy (Novocontrol Concept 81, Germany). For frequencies 10^-2^ Hz to 10^6^ Hz alpha analyzer by Novocontrol, while for frequencies 10^6^ – 3*10^9^ Hz, Keysight 4991E analyzer was used. Temperature 25°C was stabilized by the Novocool system (liquid nitrogen based) with accuracy 0.1°C. The capacitors for both, low and high frequency measurements were equipped with upper electrode with d=5 mm diameter. The height was set by a teflon spacer (h=0.13 mm). The upper electrode position was shown in the Figure 1.

**Figure 1.**
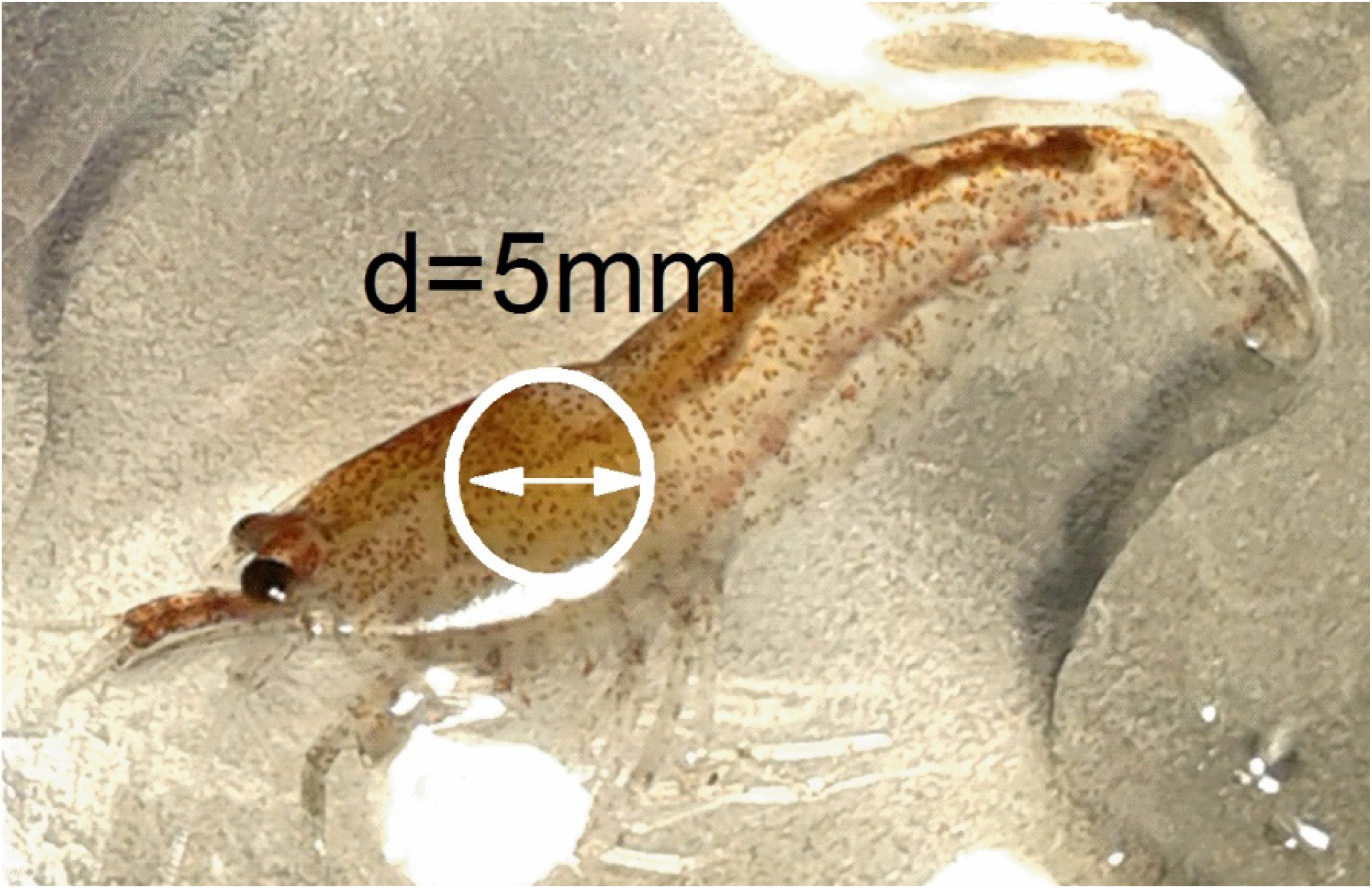
The placement of upper electrode on the body of shrimp.

Two groups i.e. control and shrimps starved by 14 days were measured by thermogravimetry method (TG) in order to obtain mass percentage of water. The TG curves were obtained by measuring shrimps after dielectric measurements in the Al_2_O_3_ crucibles with use of Netzsch STA F3 Jupiter. The heating rate 2 K/min and protective gas flow (argon 100 mL/min) were used during experiment.

## 3. Results and Discussion

Dielectric measurements were performed on five individuals from each studied group (control, starved etc.). Dielectric spectra are specific for every individual due to the differences in body weight or initial body composition. However in this work differences between studied physiological states need to be find. In order to address differences between states, dielectric spectra of organisms from every group have been averaged. Two parameters in the frequency range 10^-2^ – 10^6^ Hz have been analyzed i.e. electric permittivity and conductivity. As it was earlier stated in the Introduction section, primary relaxation process can be hardly visible in the dielectric spectrum of biological material due to the electrode polarization in the low frequencies region. On the other hand, high conductivity of measured samples covers any secondary processes in the dielectric losses. In order to observe β-relaxation in the dielectric spectrum of shrimps, differential form of Kramers-Krönig equations [12] have been applied to obtain dielectric losses from the real part of permittivity.

### 3.1 Electrode polarization

This significant decrease of σ’ in low frequencies (below 10 Hz) is caused by the electrode polarization which is related to the concentration and mobility of free ions on the surface of measured sample [13,14]. At certain limit frequency of electric field the time is sufficient for the creation of double ionic layer on the electrodes surfaces by the free ions [15]. Although in most cases electrode polarization is a parasitic process that not allows to observe true dielectric parameters of sample, in this comparative study it might be useful. The most interesting result that was found is the dependence of onset frequency at which electrode polarization occurs on the time of starvation. In the Figure 2 conductivity plots were shown for every studied physiological group (the values of conductivity are not absolute, curves on the diagram are shifted in order to make cross-over visible). In the conductivity behavior there is characteristic cross-over in the low frequency region (0.1-10 Hz), which is also the onset frequency of electrode polarization phenomenon.

**Figure 2.**
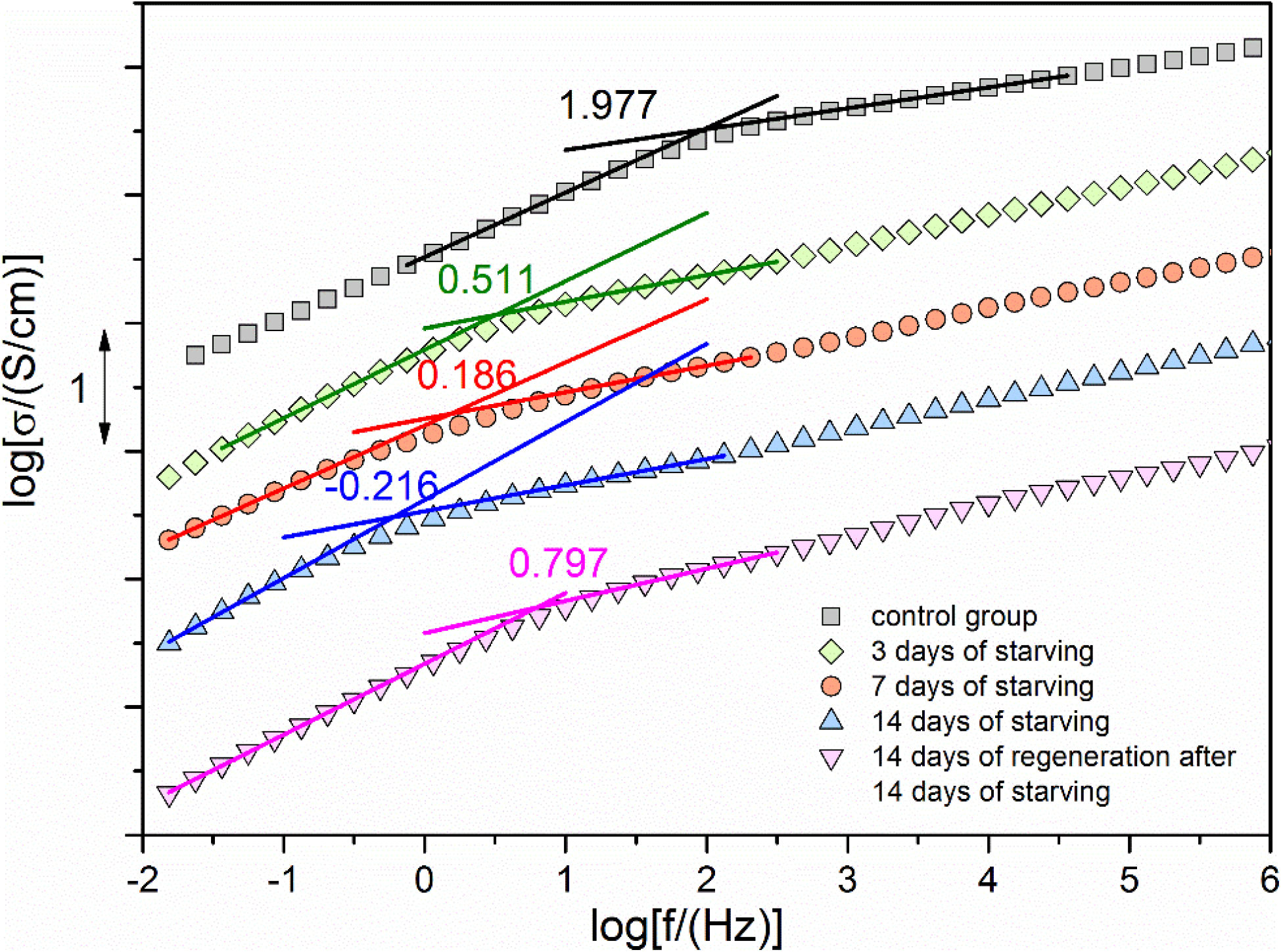
The conductivity plots for all the studied groups of shrimps. The values of conductivity are relative, the onset frequencies of electrode polarization have been shown.

The described electrode polarization onset frequencies can be used for showing reaction of shrimps to starvation. By plotting log[σ] vs time, one can obtain diagram of progress of changes occurring in a shrimp organism during starvation (see Figure 3 for details). As one can see the crossing (limit) frequency is shifting towards lower values rapidly during the starvation process. After 3 days from the beginning of experiment changes in conductivity reach the 60% of maximal value obtained after 2 weeks of experiment. However the question arises, what is the origin of observed changes in conductivity and electrode polarization behavior during the experiment. To address this question we will refer to ultrastructural changes occurring in the digestive system during starvation, which were earlier described [9]. The changes in low frequencies are related to the changes in ionic transport through the body of a shrimp. Therefore, in order to find correlations between charge transport and ultrastructural changes, analysis of changes in the cytoplasm would be most appropriate. In the cytoplasm authors have observed formation of autophagosomes. In the 4-day starved shrimps, autophagosomes were identified in the 20-25% of cells, while in 7-day starved shrimps they were identified in 40% of cells. It was also found that the number of apoptotic cells is much greater. During the apoptosis cytoplasm becomes slightly depleted of water and leads to the cell shrinking [16]. In consequence the viscosity of cytoplasm is rising and the charge transport through the cells is more difficult. Therefore, the longer the starvation period, the slower are ions and limit frequency of electrode polarization is shifting towards lower values of frequencies. In the Figure 3 the exponential decrease of onset frequency logarithm in time has been shown. Although the fact that the shrimp can survive 3 weeks on average without food, the survival mechanisms are triggered almost instantly. After 3 days of starvation physical properties are altered in a degree which is close to the 2 week progress.

**Figure 3.**
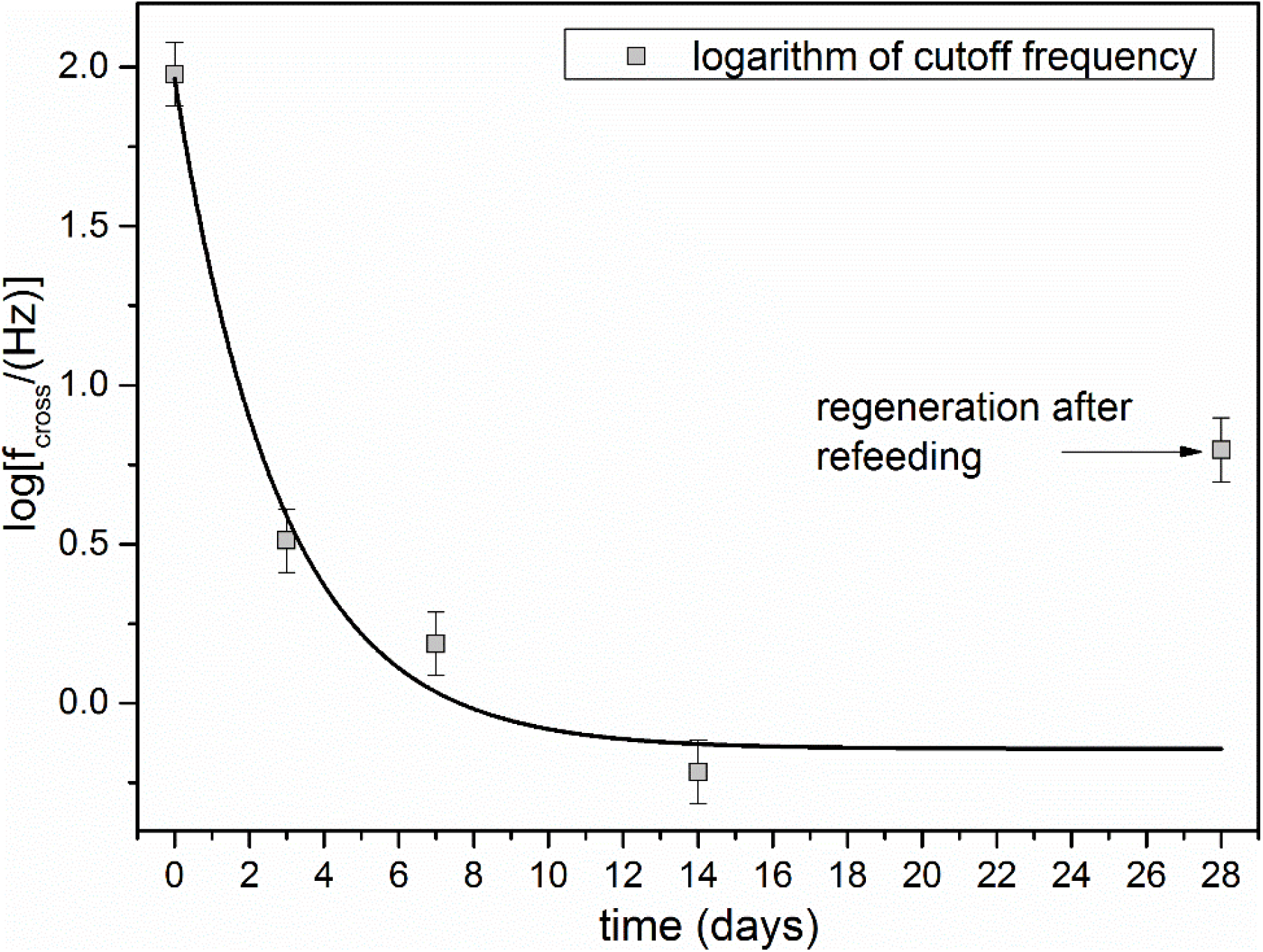
The logarithm of limit frequency of electrode polarization as a function of starvation time of shrimps.

The dielectric properties of shrimps which undergo 2 weeks of regeneration after 2 weeks of starvation are different than in control group. It can be concluded that the composition of cytoplasm of cells of organisms that survived stress induced by starvation has been changed permanently.

In order to measure the percentage of water in the body composition thermogravimetric measurements of shrimps from control group and from 14 day starved group have been performed. The observed mass loss indicates that globally, shrimps from control group consist of 52 %, while the starved shrimps consist of 68% of water. Therefore the water to other macro substances ratio has been significantly changed during starvation. Although there is an increase of water to other components ratio, during the apoptosis there is an effect of cell shrinking because of a water depletion. On the other hand apoptosis was found only in the cells of digestive system during starvation.

The absolute values of conductivity were compared for a 100 Hz frequency (above electrode polarization zone). There are no statistically significant differences in the absolute values of conductivity at the studied frequency. The measured conductivity stays in the range from 8.2*10^-7^ to 2.3*10^-5^ S/cm for all studied shrimps at f=100 Hz.

### 3.2 Relaxations

The high conductivity of samples is the reason why in the dielectric loss spectra dielectric processes i.e. dielectric peaks are not visible. In order to allow dielectric loss analysis, real part of electric permittivity can be transformed to the imaginary part by performing Krammers-Krönig transformation. In this work differential approximated form of Kramers-Krönig relations have been used i.e. 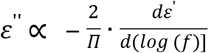.

In the Figure 4 the two relaxation processes have been revealed. The slowest process i.e. α-relaxation can be observed in the frequency range 0.1-1 Hz. In the literature, α-relaxation in tissues and cell suspensions has been attributed to the counterion polarization on the cells’ surfaces [17]. This process should be correlated to the diffusion of ions in the cytosol to the cell membrane, thus it has to be related to the viscosity of cytosol. It is clear that when the organism is in the starvation period, the composition of cytosol is changing [18–20]. In the work [9] authors have shown that in the cells of digestive system of shrimp there are active processes such as autophagy and apoptosis. These processes are aimed into recovering energy from different cell organella. The organella are being merged in the autophagosomes which can be observed just after few days of starvation. The most striking in the Figure 4 is the shift of α-relaxation towards lower frequencies during the starvation. This is clearly visible when one compare spectra for control and the shrimps after 3 days of experiment. Shifting of α-relaxation makes β-relaxation visible as an excess wing of α-relaxation. The separation of β-process is clearly seen in the spectra of shrimps after 3 days of starvation and in the group regenerated by 14 days. Although there are no clear peaks visible in the spectra, the dielectric loss curves from the 3-day and regenerated groups have been fitted by 3 Havriliak-Negami dielectric functions [21–23] which are given by the equation:

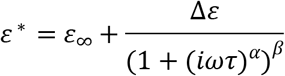

**Figure 4.**
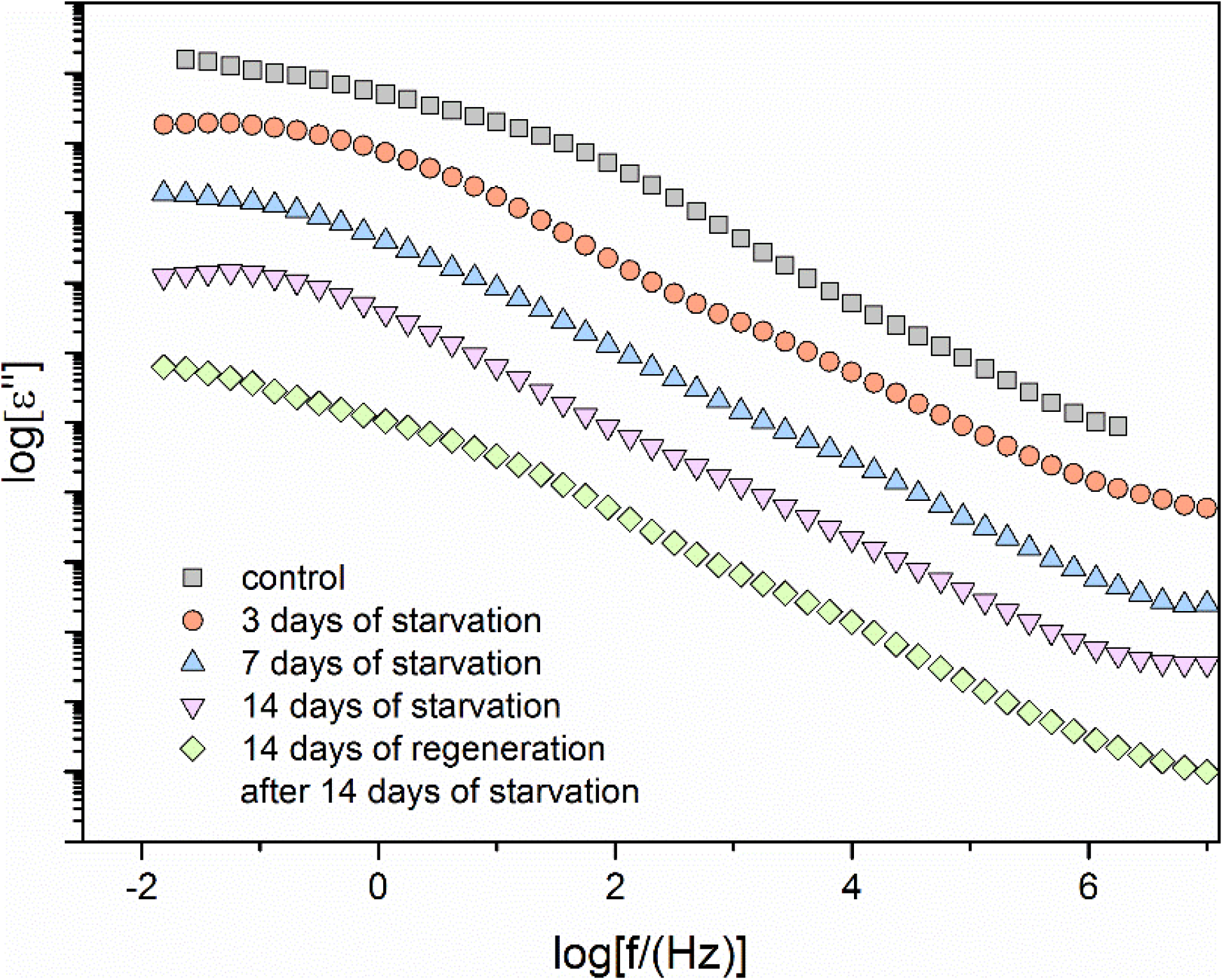
The dielectric loss spectra derived from Krammers-Krönig relations for all studied shrimp groups.

Where ε_∞_ is epsilon infinity, Δε is dielectric strength, α and β are shape parameters and τ is HN relaxation time which in case of β=1 is a characteristic relaxation time τ_max_. First HN function describes the contribution from the electrode polarization effects, second HN function describes α-process and third one describes β-process. The β shape parameter has been fixed to 1, which turns HN function into Cole-Cole function. In this case τ parameter in the fit is equal to τ_max_ of the dispersion curve and a shape parameter indicates the broadening of the curves. The peaks are symmetric. In the table 1, the parameters for two fitted curves have been shown.

**Table 1.**
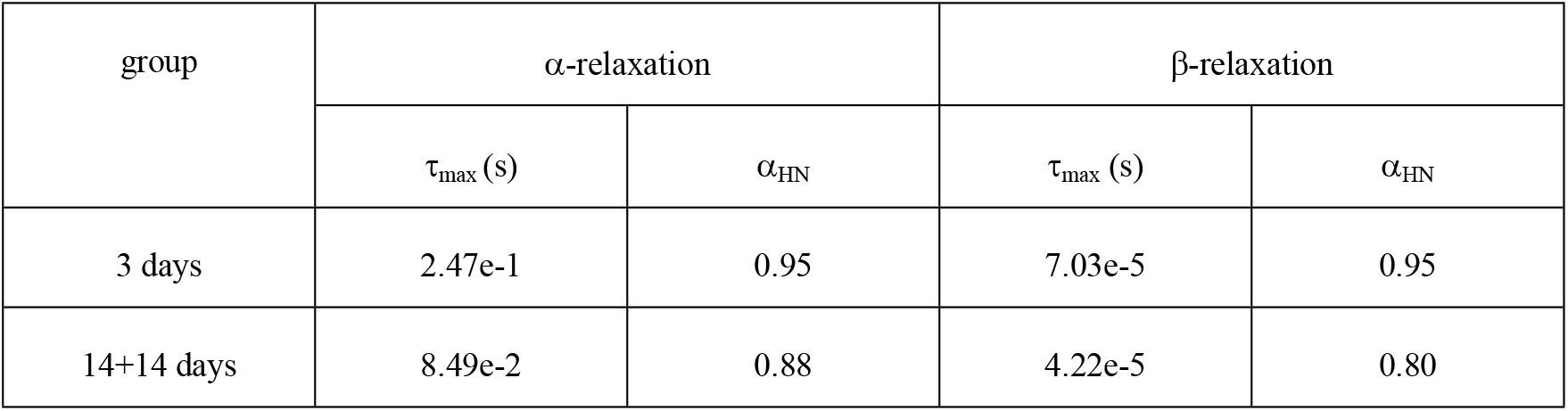
HN parameters for α and β processes have been gathered. Unfortunately the values are only approximate due to the significant overlapping of all processes in the spectra. The peaks are not visible in the spectra, only the right side of the peak.

| group | α-relaxation | | β-relaxation | |
| --- | --- | --- | --- | --- |
| | $\tau_{\max}$ (s) | $\alpha_{\text{HN}}$ | $\tau_{\max}$ (s) | $\alpha_{\text{HN}}$ |
| 3 days | 2.47e-1 | 0.95 | 7.03e-5 | 0.95 |
| 14+14 days | 8.49e-2 | 0.88 | 4.22e-5 | 0.80 |

Although the processes occurring in the shrimp bodies are highly overlapped, by performing careful fitting procedure with use of superposition of 3 HN functions, approximate parameters such as shape parameter and relaxation time have been acquired. The relaxation processes in case of 3 days starved shrimp are closer to the Debye processes. They are slightly broadened with α_HN_ parameter equal to 0.95 in both cases. In case of shrimps starved by 14 days and then regenerated by the next 14 days the processes are broader, which means broader distribution of relaxation times (α_HN_=0.88 for α-process and α_HN_=0.80 for β-process). Such situation might be related to the higher heterogeneity of the cells. In case of a-relaxation, the difference in relaxation time is noticeable. The process is significantly faster in the case of regenerated shrimps. This stays in agreement with the conductivity analysis, where limit frequency of electrode polarization is lower in starved organisms. This process clearly depends on cytoplasm viscosity. On the other hand, β-process relaxation time is similar in both cases, 7e-5 and 4e-5 s in 3-day group and 14+14 group respectively. The most pronounced β-relaxation peak has been observed in 3-day group and in the group after regeneration. This suggest the possible origin of this process. Every organism has reserve material mainly in the form of lipid droplets [19,20,24]. Such lipid droplets have been found in the hepatopancreas of *Neocaridina davidi* shrimps [25]. It can be noticed that the β-relaxation which is visible as excess wing in the spectra becomes suppressed when the starving period becomes elongated. However in the regenerated group β-relaxation is once again clearly visible. It can be connected with the fact that during starvation organisms use the reserved material in the form of lipid droplets. Re-feeding shrimps that were starved by 14 days, causes significant increase of reserve material in the extent amount which can be connected for the fact that organism is trying to prepare itself to the next foodless period [8]. Therefore molecular mobility of lipids can be related to the β-relaxation observed in the spectra of shrimps.

## 4. Conclusions

In summary, dielectric spectroscopy found to be relevant technique for monitoring the starvation process in shrimps. The progress of this process can be monitored by observing the limit frequency of electrode polarization which can be correlated to the change of ions mobility in the shrimp body. Along with the change of electrode polarization limit frequency, the α-relaxation which can be connected with counter-ion polarization of cell membranes, is shifting towards lower frequencies (becomes slower). Therefore these findings indicate that the viscosity of cytoplasm of cells increases during starvation. This can be related to the apoptosis (programmable cell death) during starvation, which is responsible for water depletion in the cells.

It was also found that there is an excess wing on the right side of α-relaxation peak which is probably a β-relaxation process hidden under the α-relaxation. The β-relaxation peak can be observed in 3 day starved shrimps, then it is suppressed for 7 day and 14 starved. It is clearly visible once again in the case of 14 days regenerated animals, while in the control group it is fully covered by the α-relaxation process. This seems to be related with the mobility of macromolecules such as lipids. The lipids are being used as the energy source during the stress induced by starvation, therefore the concentration of lipids can be coupled to the β-relaxation dielectric strength.

The last interesting finding is the large difference in the ionic mobility between control group and shrimps regenerated after starvation. The dielectric parameters of regenerated shrimps do not back to their initial values, which indicates that there are permanent differences in the body composition between control group and the shrimps that survived food shortage period during their lives.

**Figure 5.**
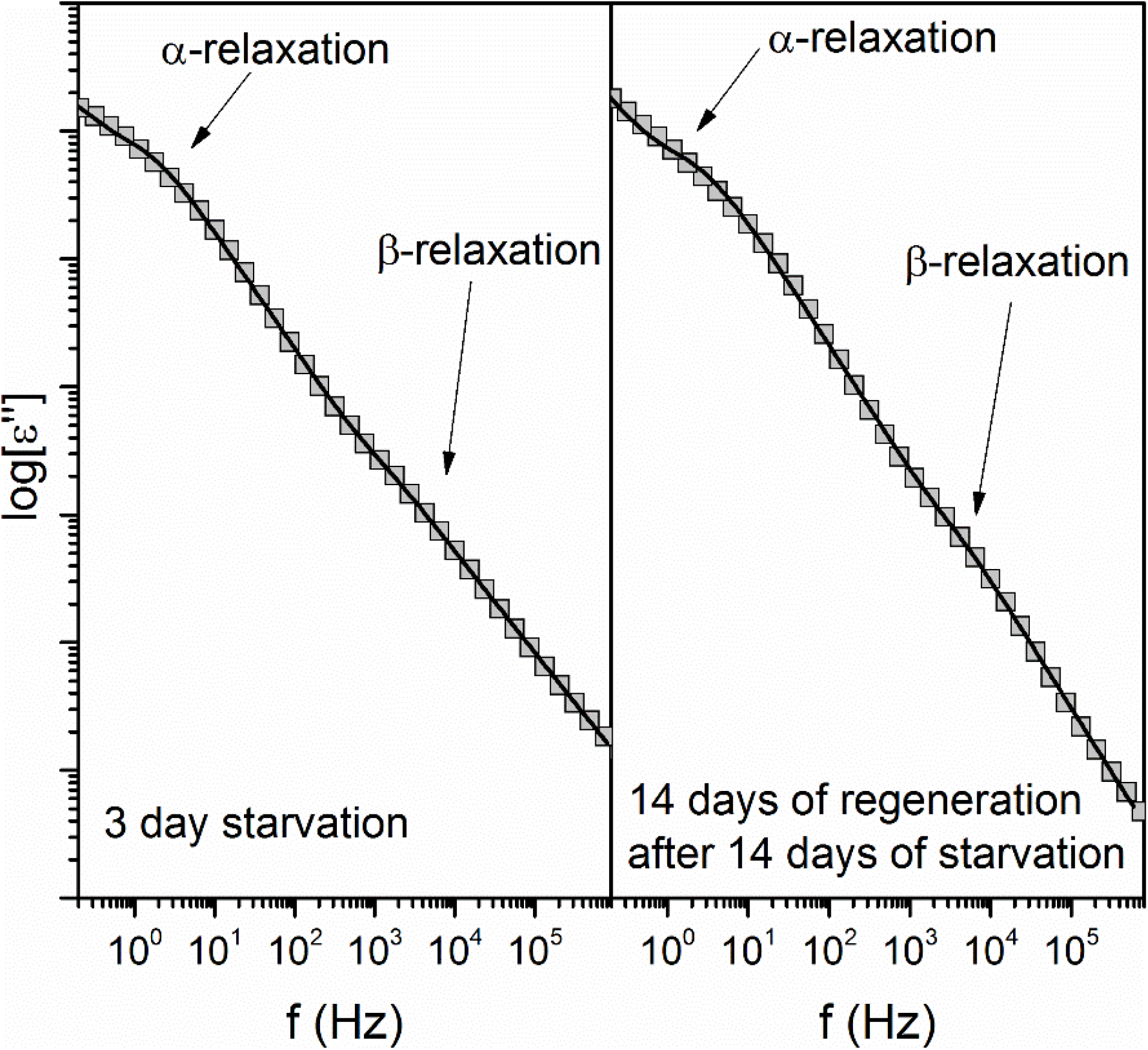
The comparison of dielectric loss spectra of 3 day starved shrimp with the one regenerated by 14 days after 14 days of starvation.

